# Conductivity of the phloem in *Mangifera indica* L.

**DOI:** 10.1101/2021.01.19.427255

**Authors:** Barceló-Anguiano Miguel, José I Hormaza, Juan M Losada

## Abstract

*Mangifera indica* is the fifth most consumed fruit worldwide, and the most important in tropical regions, but its anatomy is quite unexplored. Previous studies examined the effect of chemicals on the xylem structure in the stems of mango, but the anatomy of the phloem has remained elusive, leaving the long distance transport of photo assimilates understudied.

In this work, we used a combination of fluorescence and electron microscopy to evaluate in detail the structure of the sieve tube elements composing the phloem tissue in the tapering branches of mango trees. We then used this information to better understand the hydraulic conductivity of the sieve tubes following current models of fluid transport in trees.

Our results revealed that the anatomy of the phloem in the stems changes from current year branches, where it was protected by pericyclic fibers, to older ones, where the lack of fibers was concomitant with laticiferous canals embedded in the phloem tissue. Callose was present in the sieve plates, but also in the walls of the phloem conduits, making them discernible from other phloem cells in fresh sections. A scaling geometry of the sieve tube elements, including the number of sieve areas and the pore size across tapering branches resulted in an exponential conductivity from current year branches to the base of the tree.

Our measurements of the phloem in mango fit with measurements of the phloem architecture in the stems of forest woody species, and imply that, despite agronomic pruning practices, the sieve conduits of the phloem scale with the tapering branches. As a result, the pipe model theory applied to the continuous tubing system of the phloem appears as a good approach to understand the “long distance” hydraulic transport of photoassimilates in fruit trees.

## INTRODUCTION

Mango (*Mangifera indica* L.) ranks fifth worldwide in production after bananas, apples, grapes and citrus, and its trade and production at the global scale has gradually increased in the last decade (according to FAO, 2020). Mango is an evergreen woody perennial fruit tree crop belonging to the family Anacardiaceae, that displays monopodial growth, and both the leaf canopy and the inflorescences develop on the youngest terminal branches (Halle *et al*., 1978). Mango trees are traditionally considered as drought-tolerant, which was previously correlated with anatomical adaptations, such as a tap-root that reaches profound soil depths, or the massive presence of laticiferous canals, which have been shown to adjust the osmotic potential under drought stress (Schaffers *et al*., 1994). Yet, there is a lack of detailed anatomical information on this important fruit crop.

Although mangoes are adapted to tropical environments, they can also be cultivated in regions with subtropical climates, being low temperature (i.e. freezing risk) the major limiting factor for its cultivation at high latitudes (Whiley, 1992; Sukhvibul *et al*., 1999; Galán Saúco, 2009; Perez *et al*., 2016, 2019). However, under those conditions in which low temperatures reduce the growth of the mango trees, agronomic practices can keep mango trees at a manageable height, resulting in short trees with compacted tapering branches, which have been recently modelled (Boudon *et al*., 2020). The geometry of individual branches has an impact on the productivity of mango trees, inferred from the perspective of the allometric relationships between biomass production (i.e. leaves) and branch vigour, by means of the different effect on the pair hydraulic traits-mechanical strength (Normand *et al*., 2008). More detailed measurements of the tissues composing the stem cross sectional areas revealed that higher phloem to xylem ratios correlated with less vigorous or dwarfing varieties (Kurian, 1992). Although counterintuitive, dwarfism is a desirable character in many tree crops, due to the compact character combined with similar productivity. While there is some information on the hydraulic traits of the xylem tissue, as in most woody perennial fruit crops, there are no reports to date on phloem structure in mango, even though this tissue is pivotal for the global productivity of the trees.

The phloem is the vascular tissue that transports photoassimilates from the source photosynthetic organs toward the sink tissues, which are the growing organs of the plant, including the roots, the meristems, fruits or the inflorescences (Esau, 1969; Heo *et al*., 2014). This continuous piping system hydraulically drives substances at long and short distances, and works according to the osmotically generated pressure flow hypothesis, proposed almost a century ago (Münch, 1930), but empirically tested in a few woody organisms recently (Savage *et al*., 2017; Losada and Holbrook, 2019; Clerx *et al*., 2020). The hydraulic function of the phloem is intimately linked with the architecture of the individual conduits composing the tube, which are connected in series, and thus follow the laws of an electric circuit with resistances to flow (the connections between tubes or sieve plates). The geometrical scaling of the phloem conduits along the trunks of trees have remained unexplored until recently, when the development of microscopy techniques have allowed detailed visualization of the micro conduits of the phloem, most remarkably the quantification of the micro pores composing the sieve plates (Mullendore *et al*., 2010). As a result, the axial geometrical scaling of the phloem conduits in the branches of trees has been revealed in forest trees (Liesche *et al*., 2016; Savage *et al*., 2017; Clerx *et al*., 2020), or in the tapering branches of shrubs (Losada and Holbrook, 2019). Scaling of the conduits goes in line with the geometry of the pores, which offer most of the resistance to sap flow in the phloem, as evaluated by the most recent models of hydraulic transport (Jensen *et al*., 2012, 2014). While growing evidence of this scaling architecture of the phloem appears to dominate woody organisms, including the variability observed in woody vines (Pace *et al*., 2015), there is an only handful of studies correlating phloem structure and function in fruit trees, despite the fact that the phloem plays a critical part on survival, fructification, and response to stress.

The aim of this work is to provide detailed information on the anatomy of the so far understudied phloem conduits in the stems of *Mangifera indica* L.. With this anatomical information, we modelled the conductivity of the phloem tissue, and compared these data with recent reports on phloem structure in other species. Altogether, these observations will set the base for future works on the effect of biotic and abiotic factors on long distance transport, and consequently on the productivity of mango trees, leading to a better agronomic management of these and other tree crops.

## MATERIALS AND METHODS

### Plant material

We used two adult mango trees from the cultivar ‘Winters’, belonging to the germplasm collection of the Institute for Mediterranean and Subtropical Horticulture ‘La Mayora’ in Málaga, south of Spain (X: 407.162,62; Y:4.068.652,56; UTM:30). Trees were 6 year old grafted onto polyembrionic Gomera-4 rootstock. Given that the trees are pruned to keep a manageable canopy, we sampled branches of increasing diameters, from the current year branches to the base of the trees, following the protocol previously used for shrubs (Losada and Holbrook, 2019). Thus, we selected three branches per tree oriented to all directions, and, in each branch, four diameters with increasing thickness: between 6-8mm, 13-15mm, 26-28mm and 44-46mm (Fig.1).

**Figure 1.**
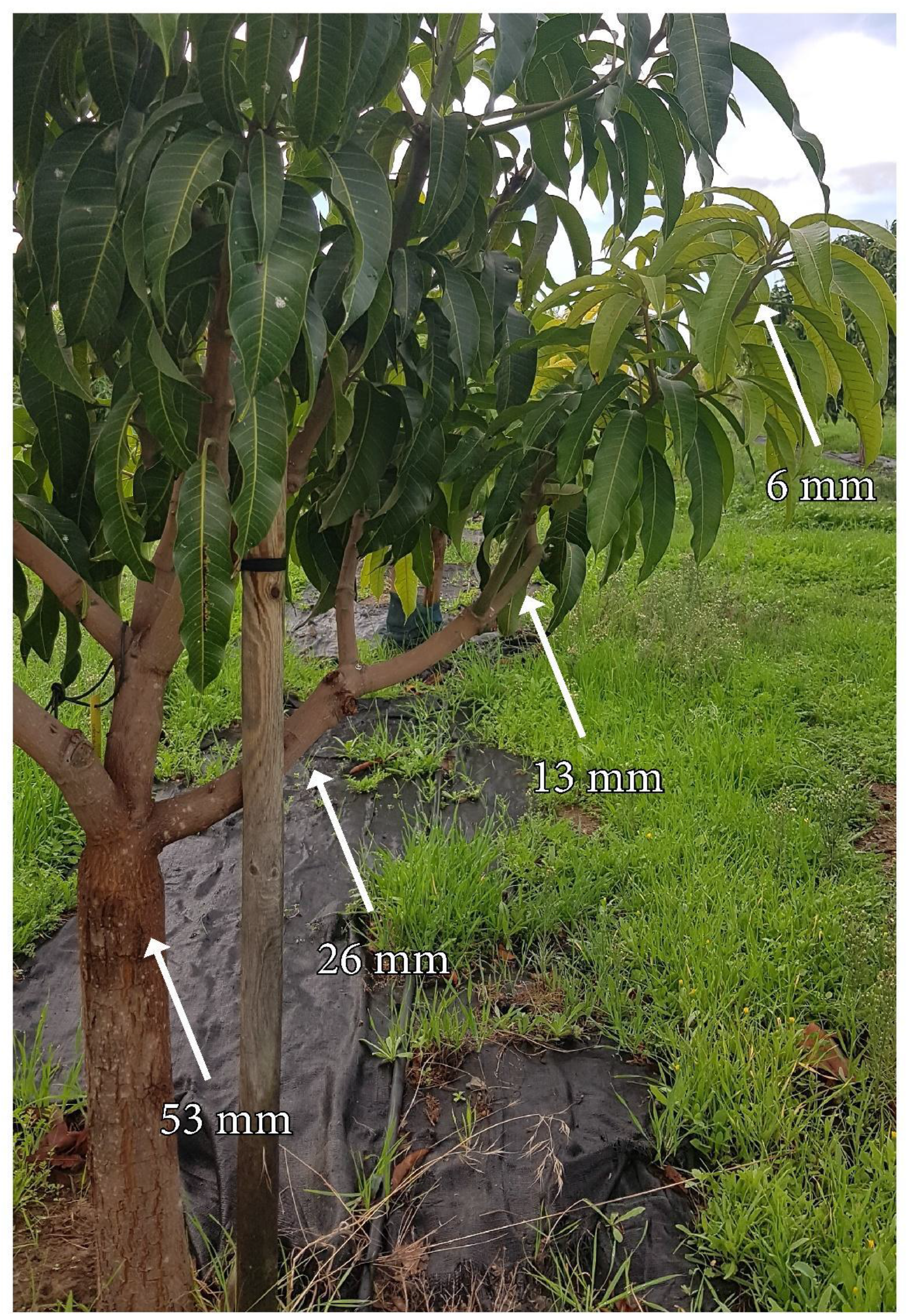
Six-year-old mango tree of the variety ‘Winters’ in the field. Arrows show the diameter of the tapering branches analysed in this study.

### Fluorescence microscopy: sieve tube geometry

Branch portions per sample point that contained the phloem tissue (i.e. the external part between the xylem and the bark), were collected with a sharp knife, and placed in 1XPBS (Phosphate – Buffered Saline), for transportation to the laboratory. Then, transverse (i.e. perpendicular to the branch length), and longitudinal (i.e. parallel to the xylem tissue) hand-sections of about 1mm thickness were obtained with a microscalpel, and mounted on slides and stained. The samples were stained either with just aniline blue (0.1% w/v in 0.1% K_3_PO_4_) (Currier and Strugger, 1956), which labels the polysaccharide callose, typically accumulated in connections between sieve tubes of the phloem –the sieve plates-, or with aniline blue counterstained with 0.1% w/v calcofluor white in 10 mM CHES buffer with 100 mM KCl (pH 10) (Hughes y McCully, 1975), which stains the cellulose of the cell walls. These sections were observed with a LEICA DM LB2 epifluorescence microscope (Leica Mycrosystems, Wetzlar, Germany), using a 405nm filter barrier, equipped with a Leica DCF310 FX camera and a LAS V4.5 software.

### Scanning electron microscopy: sieve plate pore size and density

Small portions of the branches from the same areas sampled above were cut and immediately submerged in liquid nitrogen to avoid hyper accumulation of callose in the sieve pores as a response to damage, and transferred to super chilled ethanol. They were then incubated overnight in Falcon tubes at −80°C, and gradually transferred to increasing temperatures of - 20°C for 24h, −4°C for 24h, and room temperature. The samples were then sectioned in ethanol at 1-2 mm using a double-edged razor blade. These sections were washed thrice in distilled water during 2-3 hours to remove the ethanol, and then incubated in a mixture of 0.1% w/v proteinase K dissolved in 50 mM Tris-HCl buffer, 1.5 mM Ca^2+^ acetate and 8% Triton X-100, pH 8.0 in a water bath at 60°C for at least 14 days, replacing the mixture every week (Mullendore *et al*., 2010). Once the cytoplasmic content was digested, sections were rinsed once in ethanol, and then twice in deionized water. These sections were incubated again in a water bath at 60°C for two days in 0.1% alpha amylase, which removes the starch accumulated in the sieve plates. After that, they were rinsed again in water to remove debris, and the water was poured prior to lyophilisation with a freeze-drier (CoolSafe 4–15L Freeze Dryers, LaboGene, Allerod, Denmark) for 24 hours. Desiccated samples were mounted on SEM studs, covered with gold-palladium using a sputter coater (QUORUM Q 150 R ES), and finally observed using a scanning electron microscopy (JEOL JSM-840).

### Image and data analysis

Using the fluorescence microphotographs, 75 tubes per sampling point and tree were evaluated to determine tube length and radius, giving a total of 600 tubes analysed. A total of 50 tubes were further measured per sampling point and tree to determine sieve plate number and individual sieve plate area (n=400). From the scanning electron photographs, measurements were taken in more than 75 pores per sampling location, including pore radius, pore density and the total estimated number of pores with a minimum of 5 sieve plates per sampling location. All images were analysed with the ImageJ-Fiji software (Schindelin *et al*., 2012).

Statistics were performed using SPSS 23.0 (SPSS Inc., Chicago, USA). All data followed a normal distribution, except pore area and density, which were log-transformed to fit normality. A one-way ANOVA test was used to observe differences between geometrical parameters between sampling locations. When data did not fit to a normal distribution, the non-parametric Kruskall-Wallis test was used.

### Conductivity of the phloem along the stems of mango

With the anatomical data obtained, we computed the conductivity of the phloem in the stems of mango, following the Hagen-Poiseuille model of laminar flow through cylinder pipes:

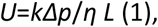

where U is the velocity of the flux, Δ*p* the differential pressure between the source and the sinks, *η* the viscosity of the fluid, and *L* the length of the tube. The first three parameters require *in vivo* measurements, and so far there are no methods available to get measurements of fluid viscosity, pressure or velocity in woody stems, with the exception of radioisotopes used to measure velocity (Orians *et al*., 2004). Thus, we focused on the evaluation of the conductivity *k*, which is greatly influenced by the variation of the geometrical parameters of the sieve tubes along the plant (see Knoblauch *et al*., 2016). For the calculation of conductivity, we used the mathematical model previously developed by Jensen *et al*. (2012, 2014), and applied to different systems such as long vines (Knoblauch *et al*., 2016), tall trees (Savage *et al*., 2017, Clerx *et al*., 2020), or shrubs (Losada and Holbrook, 2019). This method establishes that the hydraulic resistance of a sieve tube is composed by the sum of two factors, the resistance of the lumen (the axial wall of the tube) plus the resistance of the sieve plate (the connection between tubes), as to:

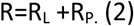

The lumen, assumed in this work as a cylinder without lateral leakage, offers a resistance that is directly proportional to the viscosity and the length of the tube, and inversely related with the fourth power of the tube radius, as to

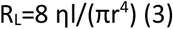

On the other hand, the sieve plate resistance is defined by the equation

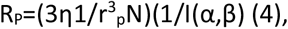

where N is the total number of pores per end tube, and I is a function of two non-dimensional parameters α and β (see Knoblauch *et al*., 2016 for details).

## RESULTS

### Anatomy of mango stems

Mango trees share a common body plan with branches that are mainly woody, except for the current year apical growth (6-8mm width), which is typically green (Fig. 1). Cross section of the current year flush displayed an extensive pith surrounded by a ring diffuse xylem; axial phloem areas around the xylem were interspersed with parenchymatous tissue; numerous laticiferous canals surrounded the phloem, and, at the most external side, they were protected by pericyclic fibres (Fig. 2A). Close up images of the cross sections of the phloem revealed the presence of callose in the walls of the sieve tube elements, which served to distinguish them from the rest of the phloem cells (Fig. 2B). In branches of wider diameters, as the pith became smaller, the xylem constituted a wider part of the cross sectional area, surrounded by a continuous phloem tissue devoid of peryciclic fibers (Fig. 2C). In those wider branches, the conduit elements of the xylem and the phloem showed longer radius and displayed a callose-rich wall (Fig. 2D). The laticiferous canals were interspersed with the sieve tubes of the phloem tissue.

**Figure 2.**
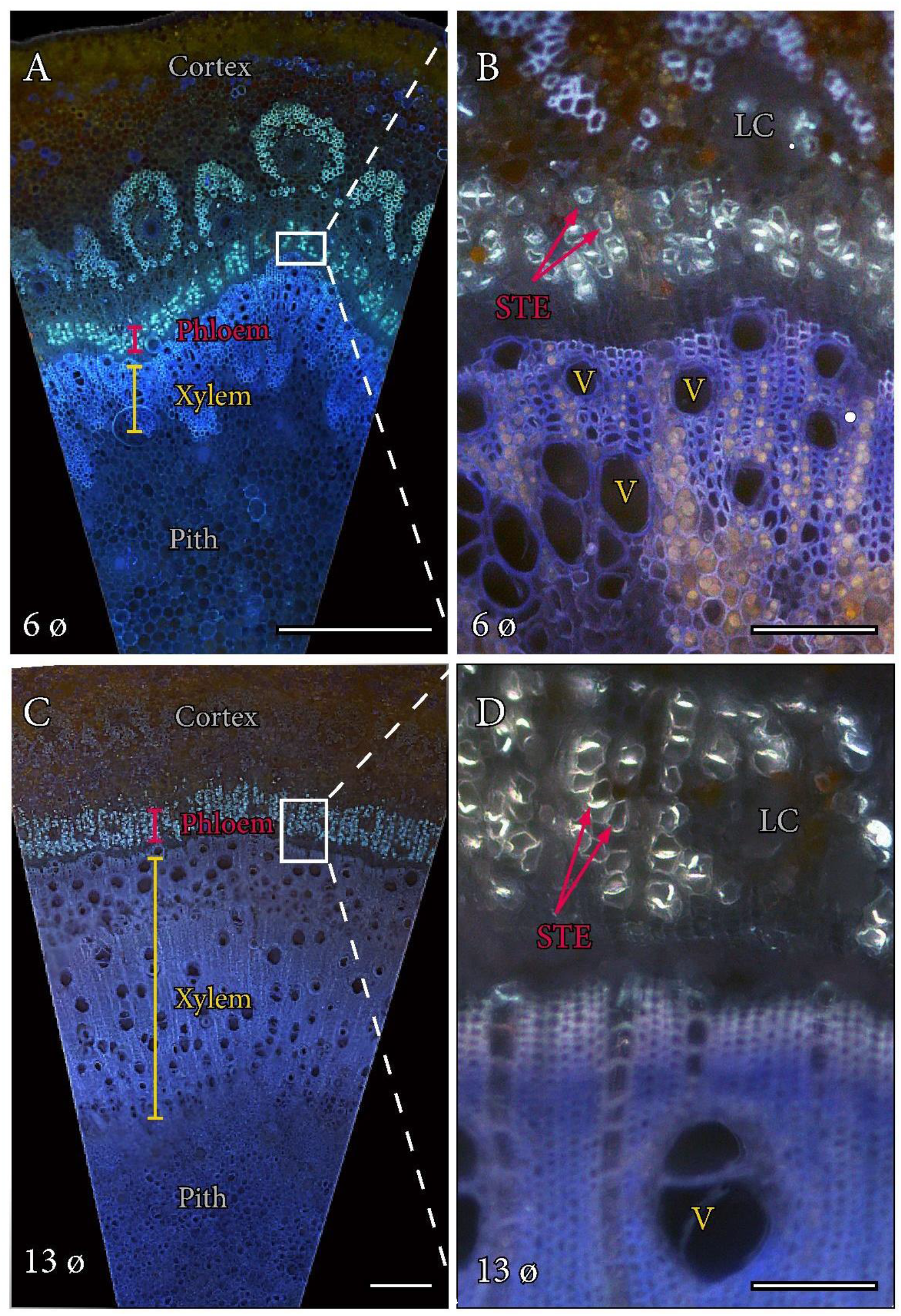
Anatomy of the stem in *Mangifera indica* ‘Winters’. A. Cross section of the 6mm stem showing the different tissues from the most external (cortex) through the phloem (red), the xylem (yellow), and the most internal (pith, white). B. Detail of the sieve tube elements of the phloem (red arrows) in cross section. C. Cross section of a 13mm diameter stem showing a larger proportion of the xylem tissue. D. Detail of the phloem, showing the wider sieve tube elements (red arrows), and the laticiferous canals embedded in the phloem tissue. Epifluorescence micrographs of cross sections of mango stems stained with aniline blue. LC, laticiferous canals; STE, sieve tube elements; V, vessels. A,C bars = 500 μm; B, D bars=100 μm.

### The phloem in the tapering branches of mango

Longitudinal views of the sieve tube elements stained with aniline blue for callose, and counterstained with calcofluor white for cellulose, revealed that sieve tubes typically associate in pairs, and sibling tubes shared numerous sieve areas in their lateral walls (Figure 3A-D). The compound sieve plates connected tubes that increased in length and width (Fig 3E), ranging from 174.6 ± 4.18 (SE) μm length in the thinnest branches, to 262.4 ± 5.99 (SE) μm in the thickest ones, following a logarithmic increase from current year branches to older ones (*r*^2^>0.91). Similarly, the values of radius length oscillated between 7.04 ± 0.11 μm (SE) in the youngest branches to 11.8 ± 0.18 (SE) μm in the older ones, further increasing logarithmically. Strikingly, the number of sieve areas per compound sieve plate, as well as their individual size, scaled logarithmically with the diameter of the branch (*r*^2^>0.97; Fig. 3C).

**Fig 3.**
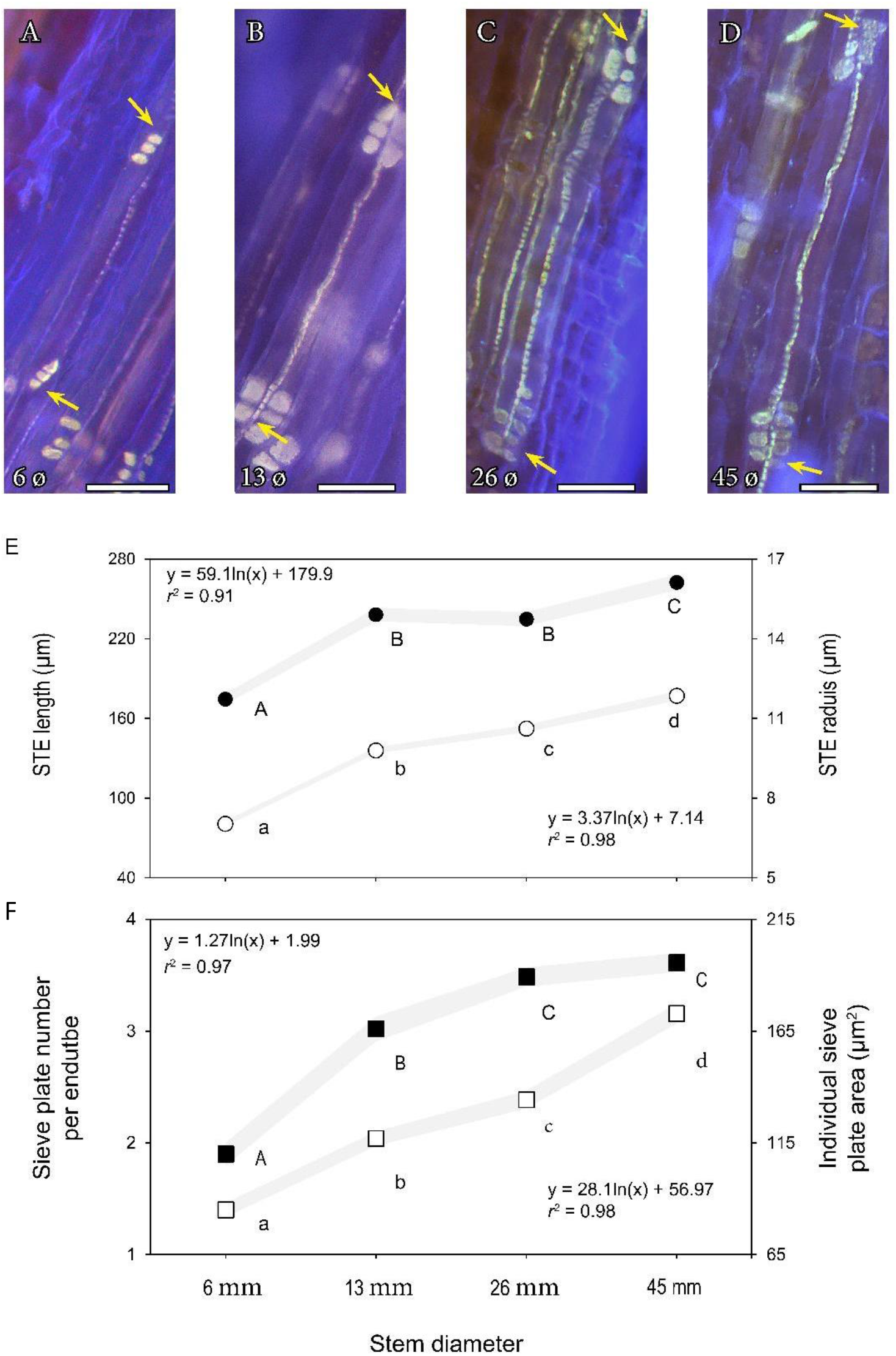
Morphology and geometry of the sieve tube elements in the stems of mango ‘Winters’. A-C Epifluorescence micrographs of sieve tube elements from branches of increasing diameters: 6 mm (A), 13 mm (B), 26 mm (C), and 45 mm (D). Yellow arrows show the compound sieve plate connections between the sieve tube elements stained with aniline blue (yellow color), counterstained with calcofluor white for cellulose (blue color). E. Sieve tube element length (solid circles) and radius (empty circles) at the four branch diameters. F. Number of sieve areas per plate connection between tubes (solid squares), and area of each plate (empty squares) at the four branch diameters. Grey lines indicate the standard error at a *p*<0.05. Letters over points display significant differences between branches following a one-way ANOVA and a Tukey test at *p*<0.05. A-D scale bars = 50 μm.

Scanning electron micrographs revealed that pore size increased gradually from the current year branch to older branches following the transport pathway (Fig. 4A-D). Quantification of the total pore number per end tube, which included the number of sieve plates as a factor, displayed a logarithmic increase with branch diameter (Fig. 4E), in line with the geometrical parameters of the tubes. The pore radius, on the other hand, further increased linearly with the diameter of the stem, roughly doubling its size from the thinnest [0.31 ± 0.01 (SE) μm] to the thickest [0.66 ± 0.01 (SE) μm] stems.

**Fig 4.**
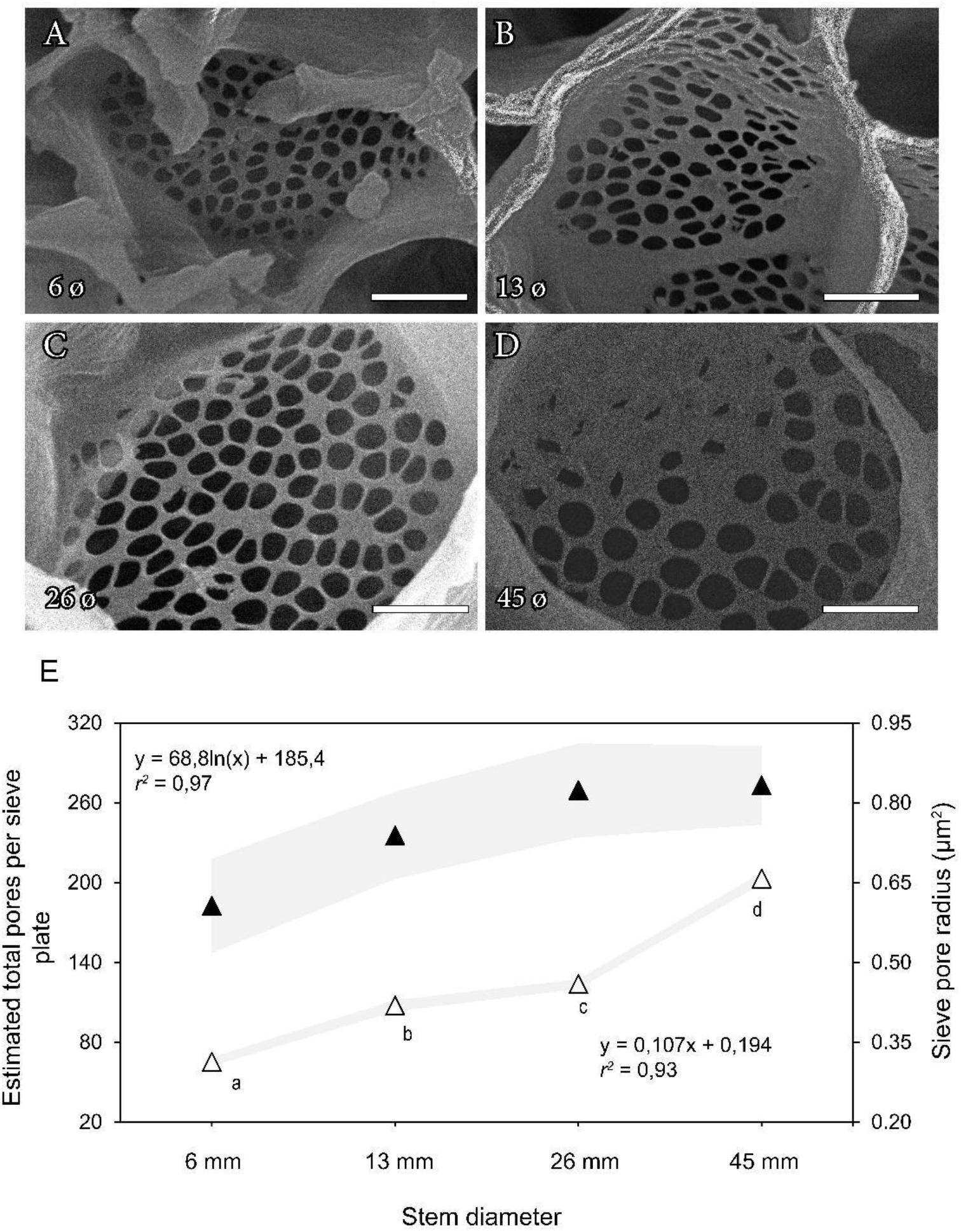
Sieve plate anatomy in mango stems. A-D. Scanning Electron Microscopy images of the sieve plates of mango “Winters” in tapering branches of 6 mm (A), 13 mm (B), 26 mm (C) and 45 mm (D). Branch diameter (bar = 4 μm). E. Estimated total pore number per compound sieve plate (solid triangles), and radius of individual pores (empty triangles) in the sieve plates of each branch diameter. Letters over symbols display significant differences in pore size at each branch diameter following a one way ANOVA and a Tukey test at *p*<0.05. A-D scale bars = 4 μm.

### Conductivity of the stems in mango

The specific conductivity of the sieve tubes in the stems of *Mangifera indica* calculated by this method oscillated between 2 and 5 μm^2^ (Fig. 5). From the youngest green branches to the thickest stems, there was an increase in the conductivity of the tubes that fitted with an exponential function, even though the trees were maintained at a height below 2m.

**Fig 5.**
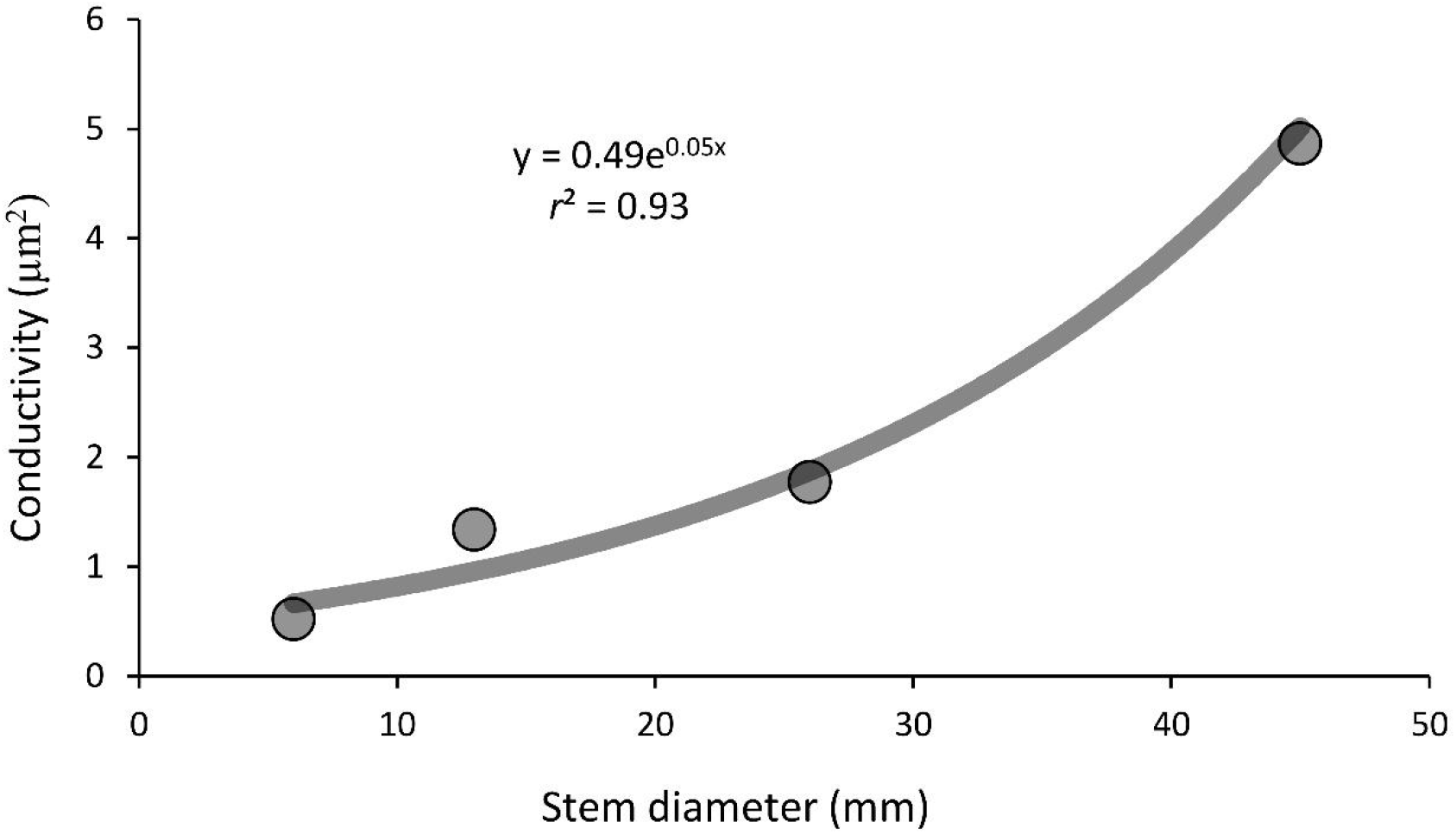
Conductivity (K) of the sieve tubes of the phloem in the stems of *Mangifera indica*.

## DISCUSSION

### CALLOSE AS A MARKER OF THE SIEVE CONDUITS OF THE SECONDARY PHLOEM

We hereby revealed that the sieve tubes of the stems in mango trees contain a basal wall content of callose, an insoluble polysaccharide that accumulates in the symplasmic connections between the sieve tube elements of the phloem, making them easily discernible from other phloem cells. Elucidating the conduit elements from other cells of the phloem is challenging in most species, yet essential to understand the general conductance of the phloem tissue, and thus the transport of photo assimilates at the whole tree level. Recently, monoclonal antibodies that attach to a branched pectin epitope present in the phloem conduits of *Beta vulgaris* leaves were applied as markers of the sieve tube walls (Torode *et al*., 2018). After that, these antibodies were used to label sieve tubes of members of the genus *Populus* (Ray and Savage, 2020), pointing to a great potential of these markers for a general use in a wide range of plant species. The technique is relatively tedious, and requires physical (i.e. using liquid nitrogen), or chemical fixation of the tissue prior to the immunolocalization, which normally leads to turgidity loss of the sieve tube elements. In order to avoid the fixation step, our work with mango showed that staining fresh sections for callose with aniline blue can be an unexpensive, fast, and relatively easy tool to identify the sieve tubes. The use of aniline blue to measure sieve tube element length in longitudinal fresh sections has been used in a number of cases (Savage *et al*., 2017; Losada and Holbrook, 2019; Clerx *et al*., 2020). However, there are few reports in which callose served to detect the sieve tubes in cross sections of woody stems, with some exceptions such as *Bombax buonopozense* and *Sterculia tragacantha* (Lawton and Canny, 1970) or the shrub *Illicium parviflorum* (Losada and Holbrook, 2019), where callose staining has been instrumental in the quantification of the cross sectional variation of the phloem tissue in the different axial positions along the stem. Yet, the technique has some inconveniences, such as the difficulty in obtaining good quality fresh sections of the phloem in a short time, which is critical to maintain the integrity of the sieve tubes. Additionally, the stiffness of the stem can be a hurdle for fresh tissue sectioning both by hand of with a sliding microtome, because the relatively flexible phloem tissue very often smashes against the rigid xylem. Despite all these inconveniences, we suggest that using aniline blue in cross sections might be a good first approach of general use to understand the structure of the phloem in tree branches.

Nevertheless, the question remains on whether callose might be present in such significant amounts in the lateral walls of the phloem conduits of other species. As a water insoluble polysaccharide, callose acts as a sealing compound, isolating cells after damage (Currier and Strugger, 1956). We can only speculate on the role of callose in the sieve tube walls, but our sections revealed that the phloem tissue of mango contains numerous laticiferous canals. Laticiferous canals pervade the majority of vegetative and reproductive tissues in most species of the family Anacardiaceae (Venning, 1948), and their role has been typically associated with defense against biotic predators, given the composition of the mucilaginous material that fills them, such as tannins, lipids and polysaccharides (Joel and Fahn, 1980). From an anatomical perspective, laticiferous canals associate with the phloem tissue, displaying positive turgor pressure due to the high concentration of osmolites putatively donated from the phloem (Pickard, 2007). The physiological role of laticiferous canals was previously studied in *Musa sp* (Milburn *et al*., 1990), *Hevea brasiliensis* (Devakumar *et al*., 1988), and *Mangifera indica* (Pongsomboom, 1991), and they seem to actively participate in the osmotic adjustment during periods of water scarcity. Callose of the sieve tubes may isolate the sieve tubes for the hydraulic function, but also provide with the mechanical resistance required to stand with positive pressures of the sap, especially during periods of drought (see Sevanto, 2014).

### CONDUCTIVITY OF THE SIEVE TUBES IN THE STEMS OF MANGO

Previous models evaluating the effect of stem diameter in current year branches revealed that they may predict leaf and branch conductivity (Normand and Lauri, 2012; Boudon *et al*., 2020). However, this refers to the conductivity of the xylem, which constitutes most of the branch mass. In addition, the geometry of the xylem vessels has been widely studied from the perspective of the scaling relationships between vessel size and branch width/height in forest trees (for example Olson and Rosell, 2012; Olson *et al*., 2020a, 2020b). In contrast, the architecture of the sieve tubes has remained understudied in many woody species, such as *Mangifera indica*. Our work showed that, in mango trees, the geometry of the individual sieve tube elements followed a direct logarithmic relationship with branch diameter, but the calculated conductivity was rather exponential, sharply increasing from the current year growth toward the base of the tree. This is because conductivity is mainly governed by the resistance of the sieve plate connections, which depends on the size and number of pores. Our detailed evaluations revealed that the sieve tube conduits contained compound sieve plates (more than one sieve area per end tube), along the branches of different widths. Compound sieve plates appear to dominate the major stems of woody species from both tropical and temperate environments (Liesche *et al*., 2016; Savage *et al*., 2017), with some exceptions that account for about 20% of tree species (Liesche *et al*., 2016). In mango (and likely in many other woody species), the number and size of the sieve areas connecting the tubes scale with the branch order/diameter, suggesting that their anatomical features are determined according to the axial position of the vascular cambium. In fact, this scaling has been recently evaluated in the stems of trees of *Quercus rubra* of different ages, which displayed compound plates (Clerx *et al*., 2020). The number of sieve areas at the base of the trunk in >5m tall trees of *Quercus* tripled those of mango, in line with their longer and wider sieve tube elements, suggesting that the age of the trunk influences the general geometry of the sieve tubes, thus facilitating transport toward the base of the tree (Clerx *et al*., 2020). The mango trees studied in this work were about 2m tall, and the geometry of the sieve tube elements falls within the values reported for *Quercus* trees of comparable heights (Clerx *et al*., 2020). Fruit trees constitute an excellent system in which to explore the effect of age on the sieve tubes, since most varieties grafted onto rootstocks maintain the genetic age of the source tree. Our sampled trees were grafted six years ago, and sieve tube anatomy displayed a scaling geometry across branches of increasing vigour. This reinforces the idea that the allometry of the vascular conduits strongly correlates with branch robustness and, thus, the hydraulic conductivity of the phloem increases in a similar fashion to that of the xylem (Normand *et al*., 2013). From a xylocentric perspective, both young (Petit and Crivellaro, 2004) and old (Olson *et al*., 2016, 2020) forest trees showed similar scaling relationships. Our quantifications of the scaling relationship between sieve pore size and diameter of the stem further reflects a conserved feature across trees of comparable ages (Liesche *et al*., 2015, 2016; Savage *et al*., 2017). We interpret this as another evidence that wider stems optimize the hydraulic conductivity of the phloem. As comparison, herbaceous vines of about 7m height, which contained simple sieve plate connections, displayed similar pore sizes, but resulted in lower conductivities than mango trees of 2m height (Knoblauch *et al*., 2016). The higher number of sieve areas per end tube in woody species might then constitute a key facilitator of the hydraulic conductivity of the phloem.

### CONCLUDING REMARKS

The unifying model that explains the structural and hydraulic function of trees, known as the pipe model theory, has typically been applied to the xylem, whose scaling depends on the environment, plant height, and other important internal and external features of the plant (Olson *et al*., 2020). In contrast, factors affecting phloem architecture are starting to be unravelled in trees, and suggest similar scaling relationships to those found in the xylem, but they deserve a careful examination in both forest and crop trees.

